# Multi-dataset Integration and Residual Connections Improve Proteome Prediction from Transcriptomes using Deep Learning

**DOI:** 10.1101/2024.07.08.602560

**Authors:** Caleb W. Cranney, Jesse G. Meyer

## Abstract

Proteomes are well known to poorly correlate with transcriptomes measured from the same sample. While connected, the complex processes that impact the relationships between transcript and protein quantities remains an open research topic. Many studies have attempted to predict proteomes from transcriptomes with limited success. Here we use publicly available data from the Clinical Proteomics Tumor Analysis Consortium to show that deep learning models designed by neural architecture search (NAS) achieve improved prediction accuracy of proteome quantities from transcriptomics. We find that this benefit is largely due to including a residual connection in the architecture that allows input information to be remembered near the end of the network. Finally, we explore which groups of transcripts are functionally important for protein prediction using model interpretation with SHAP.

## Introduction

The central dogma of biology posits a linear flow of information from genetic encoding to mRNA transcripts to functional proteins. However, this seemingly straightforward relationship belies a more intricate reality, where the interactions between omic layers are multifaceted and complex. Elucidating these relationships is crucial for understanding biological systems in both healthy and diseased states. By disentangling these interactions, researchers can identify markers and patterns specific to complex diseases, ultimately enabling the development of targeted treatments.

Among the multi-omic data type relationships, the connection between transcripts and their corresponding proteins is particularly enigmatic. Despite being quantifiable and having a direct derivative relationship, their relative quantities often exhibit only weak correlations [1], [2]. Research has identified several possible causes for this discrepancy, including alternate rates of protein generation and decay [3], [4], [5], varying reactions to environmental stimuli [6], [7], or simply systematic experimentation error bias [8], [9]. Recent works have even found that the most predictive transcripts of a protein include those that are involved in protein-protein interactions [10]. While predicting protein quantity through direct transcript-to-protein correlation remains elusive, using contextual transcript information may reveal more complex proteomic-transcriptomic relationships that could be leveraged to predict one from the other. Moreover, predicting proteins from transcripts could reduce the time and financial costs associated with future studies, as transcript quantification is generally easier to accomplish than protein quantification [11].

The National Cancer Institute’s (NCI) Clinical Proteomic Tumor Analysis Consortium (CPTAC) provides a valuable resource for exploring this issue, offering multi-omic datasets that enable research into healthy and cancerous disease states [12]. Specifically, CPTAC datasets comprise transcriptomic and proteomic data (among others) from various cancer types and adjacent healthy tissue, which makes this data useful for detection of inter-omic relationships. Several research collaborations have occurred to use this data to study RNA-protein relationships. The 2017 NCI-CPTAC DREAM Proteogenomics challenge, a sub challenge of which aimed to predict protein quantities from transcript abundance, is a notable example [13]. Contestants used a variety of methods, including random forest regression, genetic models, spline regression, linear regression, and elastic net methods, as well as ensemble combinations [14]. Using a test set only from breast cancer and ovarian cancer, the winning model achieved a pearson correlation of 0.41 and 0.47 between the true and predicted protein quantities, respectively [15]. Notably, deep learning neural networks were not strongly represented in the challenge results.

Since the close of the challenge, research into deep learning has expanded significantly, with a focus on replicating human behaviors like computer vision and natural language processing (NLP). However, the underlying principles of deep learning are equally applicable to biological data [16], [17], [18]. The key challenge lies in developing a model architecture that best fits a given problem. One underutilized strategy for deep learning model design is the NAS [19]. NAS serves a function like hyperparameter tuning, in that a range of values for specific hyperparameters are evaluated to obtain the optimal configurations. In the case of NAS, the concept is expanded to include model architecture in addition to hyperparameters, designing optimal and unique model architectures in an automated fashion. While NAS has been applied to genomic data [20], its application to predicting proteomic data from transcriptomic data remains unexplored.

We previously showed how machine learning can accurately predict the metabolome from the proteome, and how model interpretation revealed important biological insights [21]. Here, we extend that work to transcript-to-protein deep learning prediction models and demonstrate that utilizing NAS improved the accuracy. Furthermore, we highlight the potential of model interpretation to identify patterns in transcript-protein relations that underpin biological processes characteristic of a disease state.

## Methods

### Data Acquisition and Preprocessing

To facilitate readability and reproducibility, the code for downloading, processing, and splitting data was developed as a multi-class in-house package. The bridge design pattern specifically was used to allow for interchangeability of data source input to expand beyond CPTAC in the future, as well as for allowing custom processing and splitting depending on the experiment. This involved writing an abstract parent data processing class, and individual child classes would utilize compartmentalized class components specific to the experiment. This was done to enable external researchers to trace data processing workflows with ease.

CPTAC data was downloaded directly from zenodo using the cptac python package [22] separately for each cancer type. Analyses were only performed using transcripts and proteins common between all datasets.

Cancer-specific datasets were normalized independently of one another. For experiments where each dataset required an identical train-validation split, this normalization was calculated on the training partition then applied to the training and validation partitions both. For the five by two cross validation experiments, it was determined that dataset-specific analyses would be enhanced by the inclusion of all other datasets in the training dataset [**Supplementary Figure 1**]. In these instances, the train-validation split was applied only to the target dataset, with a split of 0.45, 0.45, and 0.1 for training, validation and testing, respectively. All non-target datasets were normalized on the entire dataset as the training partition, ignoring validation or testing partitioning entirely. In a standard data partition, each dataset had a split of 0.8, 0.1 and 0.1 for training, validation and testing, respectively.

The final dataset contained data for breast cancer (BRCA), kidney cancer (CCRCC), colon cancer (COAD), brain cancer (GBM), squamous cell cancer (HNSCC), lung cancer (LSCC and LUAD), ovarian cancer (OV), and pancreatic cancer (PDAC) as well as adjacent healthy tissue. 59286 transcripts and 7822 proteins were shared across all datasets with a combined number of 1256 samples.

### NAS

The search space used in this study chose a model architecture consisting of three segments, or blocks, of sub-architectures. These blocks could vary in number of neurons, number of layers, activation functions between layers, intra-block residual connections, dropout rates, or be removed entirely to simplify the network. It was determined that while mRNA to encoded protein quantities are not directly correlated, the quantity of one likely has a strong impact on the quantity of the other. Thus, the search space also included a residual connection inserting mRNA input quantities for proteins being predicted right before the final output layer of the network.

The NAS workflow was based on the “Multi-Objective NAS with Ax” workflow tutorial on the official pytorch website [23], utilizing Meta’s Ax package to do so. The process includes designing a search space as a separate python script that accepts variables that dictate the model structure, setting up a torchx runner and scheduler for submitting model training scripts concurrently, and defining optimization requirement configurations. Ax uses Bayesian optimization to evaluate and compare model configurations and their predictive accuracy, highlighting the impact specific architecture decisions have on the final loss.

### Model Evaluation and Comparison

Losses between predicted and true outputs were calculated using mean-squared error. The dummy regressor identified the mean of the true output data and used it as the prediction of all data points. The random forest regressor was run with log2 max feature and 50 node max depth limitations, as the number of inputs and outputs in creating a forest of full trees would otherwise require upwards of years to calculate. The manually designed model consisted of two hidden layers with output sizes of 12k, and 10k, respectively. Hidden layers employed batch normalization, a dropout rate of 0.6, and used the leaky ReLU activation function with a negative slope of 0.05 [**Supplementary Figure 2**]. The NAS-optimized model consisted of three blocks of layers, followed by a single output layer. The first block consisted of a single neural layer with 319 neurons, a sigmoid activation function, a dropout rate of 0.52. The second block consisted of three neural layers with 508 neurons, a sigmoid activation function, a dropout rate of 0.69, and a residual connection skipping the middle layer. The third block consisted of a single neural layer with 7822 neurons (the output size), a tanh activation function, and a dropout rate of 0.9. The optimal model also sported a batch size of 128 and a learning rate of 1e-4.

### Model Interpretation Using SHAP Values

SHAP values [24] were calculated using all available samples from CPTAC used in training and validation of the optimal model, which included the direct transcript residual connection. SHAP values for the top 13 accurately predicted proteins across cancers using the NAS optimized model were extracted and graphed independently, specifically CAVIN1, FERMT2, FLNA, HCLS1, TK3, MCM3, MCM4, MCM6, P4HB, PTPN6, SMC2, STAT1, and VCL. MMP14 was included as a candidate because of its role in cancer regulation. SHAP values for targeted proteins were extracted and analyzed independently of the raw SHAP outputs for memory efficiency purposes.

Direct transcripts were determined by directly matching gene names to the list of predicted proteins. A cutoff of –15 mean absolute SHAP value was chosen to separate correlated and uncorrelated transcripts.

Transcripts that had an absolute mean SHAP value above –15 in at least 70% of the chosen proteins were determined to belong to the correlated category. Transcripts that had a null or 0 SHAP value were excluded from categorization.

A SHAP analysis was also run on 3 variations of the optimal model, specifically models with no input residual connection, a correlated transcript residual connection, and an uncorrelated transcript residual connection. To accommodate residual connections of varying lengths, the output size of the third block and the input size of the final output layer were altered to match the size of the residual connection for these adjustments.

### Code Availability

CPTAC data was downloaded using the cptac python package [22]. All models were developed using Pytorch [25] and Pytorch Lightning [26]. NAS was implemented with Meta’s Adaptive Experimentation Platform (Ax) [27]. NAS evaluation metrics were tracked with tensorboardX [28]. Scikit-learn was used to perform train-validation splits and several non-neural net regression models [29]. Pandas was used to load, process and save CPTAC data [30]. Numpy was used to perform various calculations [31]. SHAP was performed with the shap python package [24].

Code for data processing and model training and evaluation can be found at https://github.com/xomicsdatascience/RnaToProteinDataModule. Classes for data processing and model generation can be found in the src directory, while scripts for running the different experiments can be found in the scripts directory.

### Large Language Model Edit

This paper was refined for human readability using Meta’s Llama 3 Large Language Model [32].

## Results

### CPTAC Model Configuration

The CPTAC dataset provides a unique opportunity to investigate the relationships between multi-omic layers, particularly in predicting the proteome from transcriptome data [**Figure 1A**]. A range of approaches were employed to achieve this, including one-to-one mapping of mRNA transcripts to proteins compared with various regressors: dummy, linear, random forest, and neural networks [**Figure 1B**]. The latter proved to be the most variable, with different architectures yielding diverse predictive accuracies. To address this, a NAS was utilized to automate the selection of optimal neural network architectures [**Figure 1C**]. The NAS approach iteratively evaluated various model architectures from a search space to identify the most effective structure for predicting protein quantities from transcriptome data [**Figures 1C, 1D**]. The optimal NAS model was benchmarked against other models, including a dummy regressor, and its performance was evaluated using a 100×2 cross-validation metric across different cancer types. The results showed that the optimal NAS model consistently outperformed all other models, including the previously optimal random forest model [**Figure 1E**].

**Figure 1:**
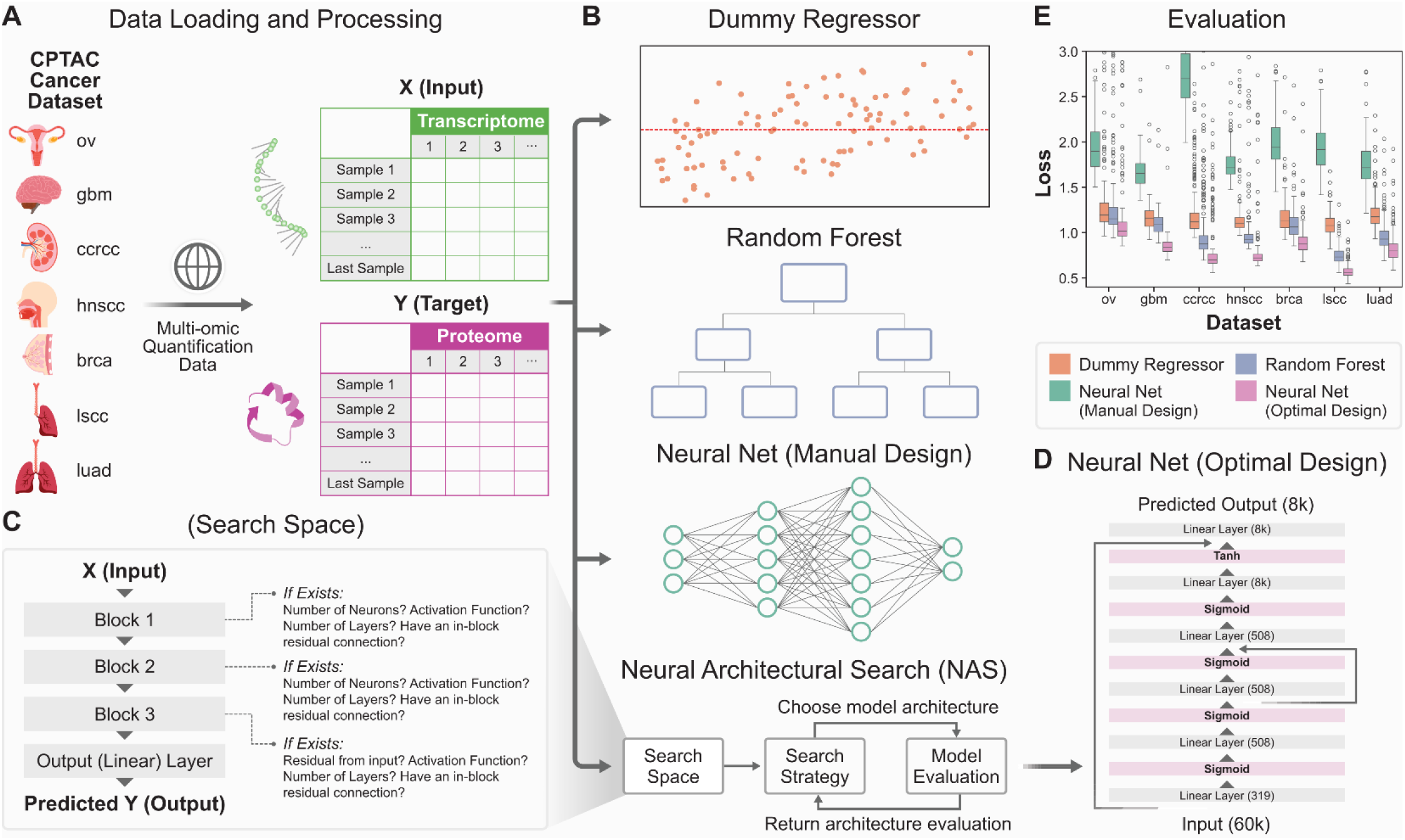
Comparison of methods for predicting the proteome from the transcriptome. **(A)** CPTAC cancer datasets were downloaded from the web and processed to match samples across omic layers. **(B)** Four primary methods were evaluated for predicting the proteome from the transcriptome. They are a dummy regressor to serve as a benchmark, a random forest regressor, a manually designed neural network, and an optimized neural network chosen via NAS. **(C)** The primary outline of the NAS search space used to identify the optimized neural network. **(D)** The model architecture of the optimized neural network. **(E)** A comparison between the four methods via 100×2 cross validation, indicating the optimized neural network outperforms all the others.

### Model Performance and Correlation Analysis

Given that the proteome and transcriptome do not reliably correlate with each other, a correlation between predicted protein quantities and actual protein quantities was calculated. An example of the RNA/protein correlation for BGN is shown in **Figure 2A**, while the true versus neural network-predicted value of the protein is shown in **Figure 2B**. R2 scores were calculated between transcripts and proteins, as well as predicted proteins to actual proteins, and compared to the benchmark transcript-to-protein correlation. These analyses revealed that the predictions from the model greatly improved the prediction for this example protein. Additionally, it was assessed whether a simple linear model could sufficiently correlate predicted proteins with actual proteins. The results revealed that the optimal NAS model outperformed both the direct and linear evaluations, highlighting its effectiveness in predicting protein quantities from transcriptome data. A global view of the R2 score distributions for each of the three relationships across all 7822 genes showed that our neural network from NAS performed best [**Figure 2C**].

**Figure 2:**
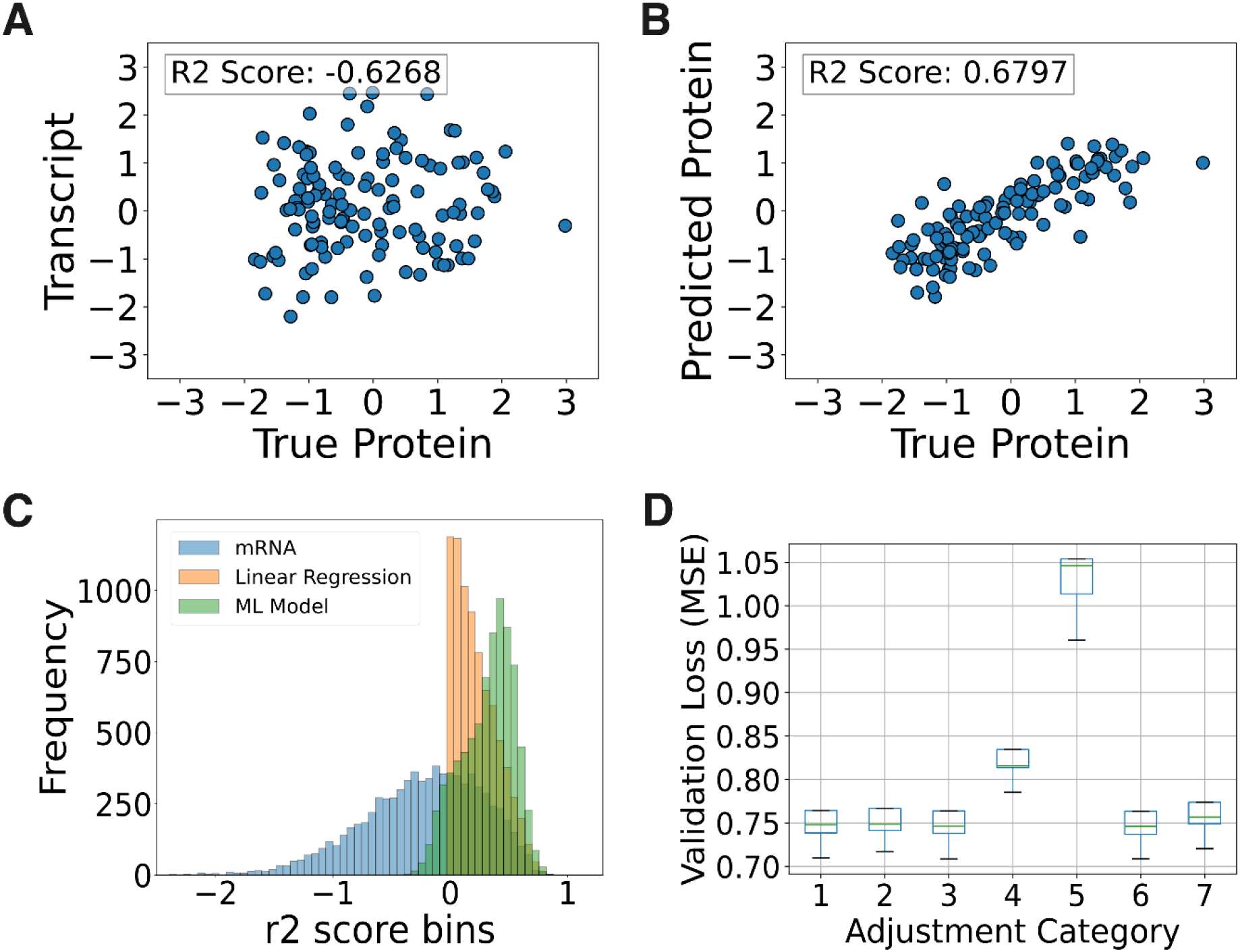
Evaluation of the impact and design of the NAS-optimized neural network. **(A)** Comparison of the normalized transcripts quantities against the matching normalized protein quantities of the validation dataset for the BGN gene. Note that the values have a negative R2 score, indicating little to no correlation. **(B)** Comparison of the normalized predicted output quantities against the matching true normalized protein quantities of the validation dataset for the BGN gene. Note the change in R2 score, indicating strong correlation. **(C)** Validation set R2 scores for every protein evaluated by the model. The different bar colors indicate which dataset the true protein quantities were compared to, specifically matching mRNA quantities (blue), the output of a simple linear regression mapping (orange), and the output of the NAS-optimized neural network (green). **(D)** An evaluation of the different unique components of the NAS-optimized neural network. Corresponding to the search space summary, the categories map to (1) removal of the first block, (2) removal of the second block, (3) removal of the residual connection in the second block, (4) changing the dropout rate of the third block, 0.9, to match that of the first block, 0.52, (5) removing the residual connection from the input to the immediately before the output layer, (6) the unchanged optimized model, and (7) using a sigmoid activation function for the third block instead of the NAS-determined tanh activation function.

Inspection of the optimal NAS model’s architecture identified key components, including a single-layer block, a three-layer block with a residual connection, and a final single-layer block with a tanh activation function [**Figure 1D**]. Notably, the mRNA residual connection played a crucial role in the model’s accuracy. To quantify the impact of each component, individual components were selectively removed or nullified to determine their impact on the output, revealing that the mRNA residual connection had the most dramatic effect on the model’s performance [**Figure 2D**]. In addition to the residual connection, multi-dataset integration across all cancers was important for improving model performance [Supplementary Figure 1].

### Biological Feature Analysis and Model Improvement

To uncover biological features underlying the transcriptome-proteome relationship, a Shapley Additive Explanation (SHAP)[24] analysis was performed to identify key transcripts correlated with protein quantities. The results showed that mRNA transcripts emphasized by the mRNA residual connection had distinctly higher average absolute SHAP values, highlighting the importance of this connection. Furthermore, a clear distinction was observed between transcripts correlated with protein quantity and those that were not, with the former exhibiting higher SHAP values. These transcripts were categorized as direct, correlated, and uncorrelated, respectively [**Figure 3, Supplementary Figure 3**].

**Figure 3:**
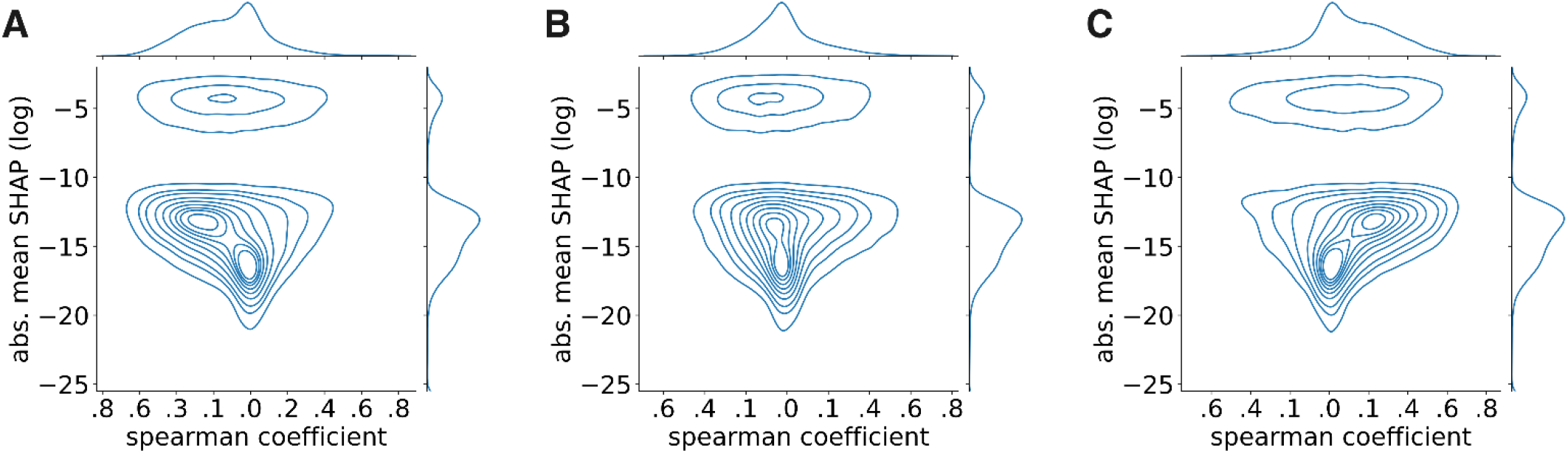
Model interpretation evaluating transcript impact on specific protein predictions. SHAP analysis was performed for several of the most well-predicted proteins using all samples in the dataset. As a correlative benchmark, the spearman coefficient between each transcript quantity and the specific protein quantity was also determined. A spearman coefficient of 0 indicates no correlation while negative or positive values indicate negative or positive correlation, respectively. A kernel density graph (KDE) was plotted for mean absolute SHAP (y axis) versus spearman coefficient (x axis) for each transcript’s relation to **(A)** MCM6, **(B)** VCL, and **(C)** FERMT2 proteins specifically. Three general patterns of transcripts appeared, namely a high-impact group, a semi-correlated mid-impact group, and an uncorrelated low-impact group.

Direct and correlated transcripts might serve similar roles in predictive performance and the residual connection using direct transcripts may bias their importance according to SHAP. Instead, incorporating correlated transcripts into the residual connection could enhance the model or alter their importance. To test this, the input residual connection was manually configured to allow for differential insertion of each transcript category. The results showed that insertion of each group into the residual connection had a different effect on model performance and their SHAP values; while uncorrelated transcripts had a marginal impact on model accuracy, direct and correlated transcripts significantly improved predictive performance. Notably, direct transcripts had the largest impact, but correlated transcripts also demonstrated a substantial effect, suggesting a relationship between these transcripts and the predicted proteins [**Figure 4**]. It is also notable that groups promoted with a residual connection had a consistent corresponding boost in SHAP values, indicating a relationship between model architecture and interpretability.

**Figure 4:**
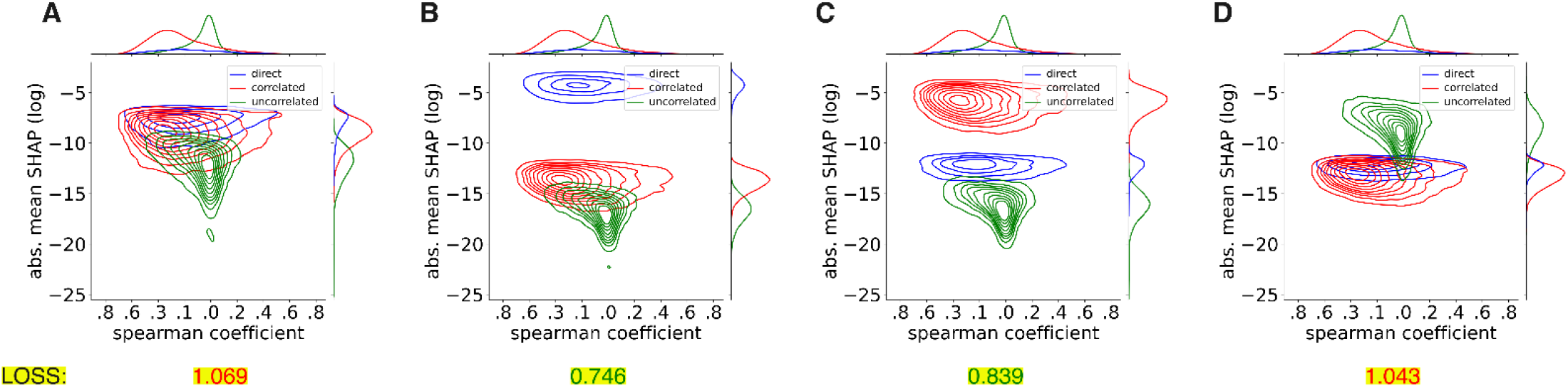
Evaluation of the input residual connection impact using several different categories of transcripts. Kernel density estimation (KDE) for the MMP14 protein. Differing categories of transcripts were applied as the input residual connection, and the resulting impact on SHAP values between each category was plotted. The training loss for each model is included as a benchmark, with red text indicating high loss values and green text indicating low loss values. (A) Evaluation of the model where the input residual connection was removed entirely. (B) Evaluation of the model where only transcripts that directly matched to target proteins were used in the residual connection. (C) Evaluation of the model where only non-direct transcripts above a log absolute mean SHAP value of –15 were included in the residual connection. (D) Evaluation of the model where only non-direct transcripts below a log absolute mean SHAP value of –15 were included in the residual connection. Notably, each time a group is used as an input residual connection, the general impact of group members exceeds transcripts outside the group. The direct transcript model ultimately outperforms all others by mean-squared error (MSE) loss metrics.

The types of transcripts in each category differed in proportion. While the direct transcripts consisted entirely of coding proteins, the correlated and uncorrelated transcripts had different mixtures of various RNA types. The correlated transcripts emphasized long non-coding RNAs and indirect protein coding mRNAs, while uncorrelated transcripts emphasized processed pseudogenes, though all three types were found in both groups [**Figure 5, Supplementary Table 1**]. Overall, the findings highlight the importance of optimizing neural network architectures and incorporating biologically relevant features, such as mRNA residual connections and correlated transcripts, to improve the prediction of protein quantities from transcriptome data.

**Figure 5:**
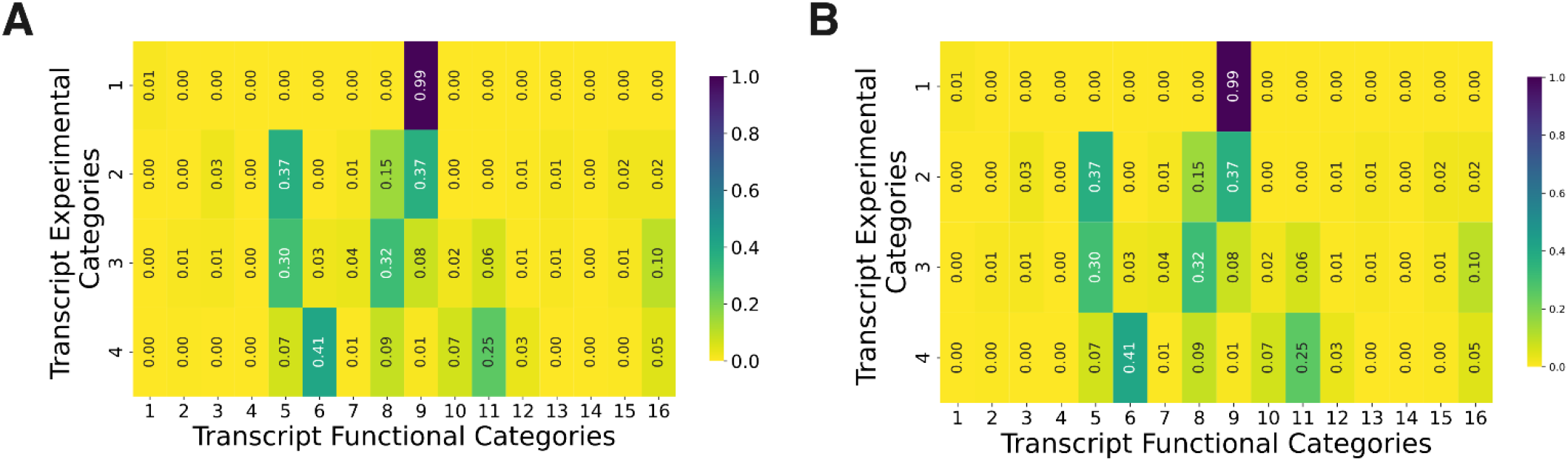
Evaluation of the functional composition of transcript categories. Transcript experimental categories are (1) direct transcripts, (2) correlated transcripts, (3) uncorrelated transcripts, and (4) transcripts in the dataset with zero quantity in the analyzed samples. Transcript functional categories are (1) ‘IG_V_gene,’ or immunoglobulin variable chain genes, (2) ‘IG_V_pseudogene,’ or inactivated immunoglobulin genes, (3) ‘TEC,’ or transcripts to be experimentally confirmed, (4) ‘TR_V_gene,’ or T-cell receptor genes, (5) ‘lncRNA,’ or long non-coding RNAs, (6) ‘miRNA,’ or microRNAs, (7) ‘miscRNA,’ or miscellaneous other RNAs, (8) ‘processed_pseudogene,’ or pseudogenes that lack introns, (9) ‘protein_coding,’ or transcripts that contain an open reading frame, (10) ‘rRNA_pseudogene,’ or non-coding ribosomal RNAs predicted to be a pseudogenes by the Ensembl pipeline, (11) ‘snRNA,’ or small nuclear RNAs, (12) ‘snoRNA,’ or small nucleolar RNA, (13) ‘transcribed_processed_pseudogene,’ or pseudogenes that lack introns and have expression indicated by locus-specific transcripts, (14) ‘transcribed_unitary_pseudogene’, or pseudogenes where the presence of locus-specific transcripts indicates expression and there is an active orthologue in another species, (15) ‘transcribed_unprocessed_pseudogene,’ or pseudogenes with introns and have expression indicated by locus-specific transcripts, and (16) ‘unprocessed_pseudogene,’ or pseudogenes with introns. (A) Heatmap of proportions of transcripts across experimental categories. (B) Heatmap of proportions of transcripts across functional categories.

## Discussion

The results of this study demonstrate the potential of NAS in optimizing deep learning models for predicting protein quantities from transcriptome data. By automating the selection of optimal neural network architectures, we were able to identify a model that consistently outperformed other approaches, including random forest and linear regression. The importance of incorporating biologically relevant features, such as mRNA residual connections and correlated transcripts, was also highlighted.

The NAS-optimized model’s ability to predict protein quantities from transcriptome data with improved accuracy has significant implications for the field of proteogenomics. By leveraging the strengths of deep learning and multi-omic data, we can gain a better understanding of the complex relationships between different omic layers. This, in turn, can lead to the identification of novel biomarkers and therapeutic targets for diseases. We expect that as larger and larger datasets become available for training; the approach we outlined here will improve this prediction further.

The SHAP analysis revealed several distinct categories of transcripts that had varying degrees of impact on protein quantity prediction. These findings suggest that the model can capture biologically meaningful patterns in the data, and that the incorporation of correlated transcripts can enhance predictive performance in future models. Although the residual connection clearly biased the group of direct transcripts, Figure 5 suggests that different classes of transcripts have different effects on protein quantities. For example, snRNA, which functions in the spliceosome to process pre-RNA, have almost no specific impact on protein quantity prediction and are found in the unimpactful, uncorrelated transcript category. Messenger RNAs generally have a high impact on predictive function. Long non-coding RNAs seem broadly more impactful for a gene’s quantity, whereas the importance of pseudogenes appears to depend on their state of processing. Only a small subset of microRNAs (1%) were important for predicting this set of proteins, supporting their role in specific biological regulation.

## Conclusion

This study demonstrates the potential of NAS and deep learning in optimizing protein quantity predictions from transcriptome data. The results highlight the importance of incorporating biologically relevant features and the need for further research into the development of more accurate and efficient models. By leveraging the strengths of these approaches, we can gain a better understanding of the complex relationships between different omic layers and make significant progress in the field of proteogenomics.

## Acknowledgements

We appreciate advice from Zijun (Frank) Zhang related to NAS and Alexandre Hutton for advice related to deep learning. Dasom Hwang helped with graphic design. This work was funded by the National Institute of General Medical Sciences (NIGMS) R35GM142502.

