## Supplemental Information for "Multi-dataset Integration and Residual Connections Improve Proteome Prediction from Transcriptomes using Deep Learning"

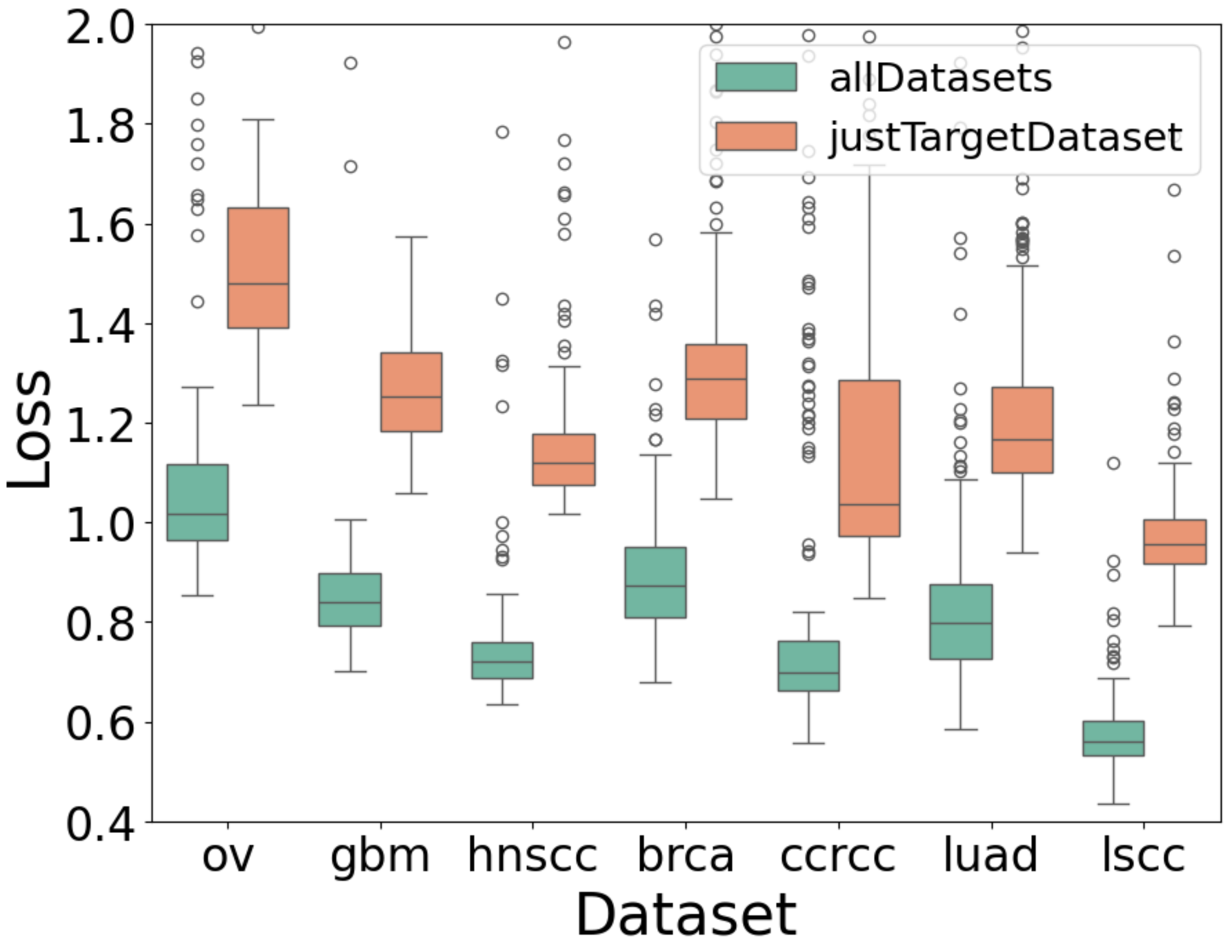


Supplemental Figure 1: **Comparing all datasets used in training vs. only target dataset use in training.** Target dataset had an 80:20 trainining:validation split for all replicate calculations. In the justTargetDataset runs, no addiional changes were made. In the allDatasets runs, the training data was supplemented with all full non-target datasets.

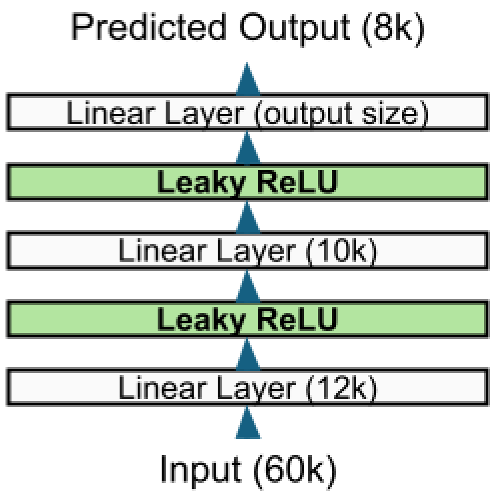


Supplemental Figure 2: **Manually designed neural network architecture.** Prior to Neural Architectural search optimization, a simple three-layer network with leaky relu activation functions was trained on the CPTAC datasets.

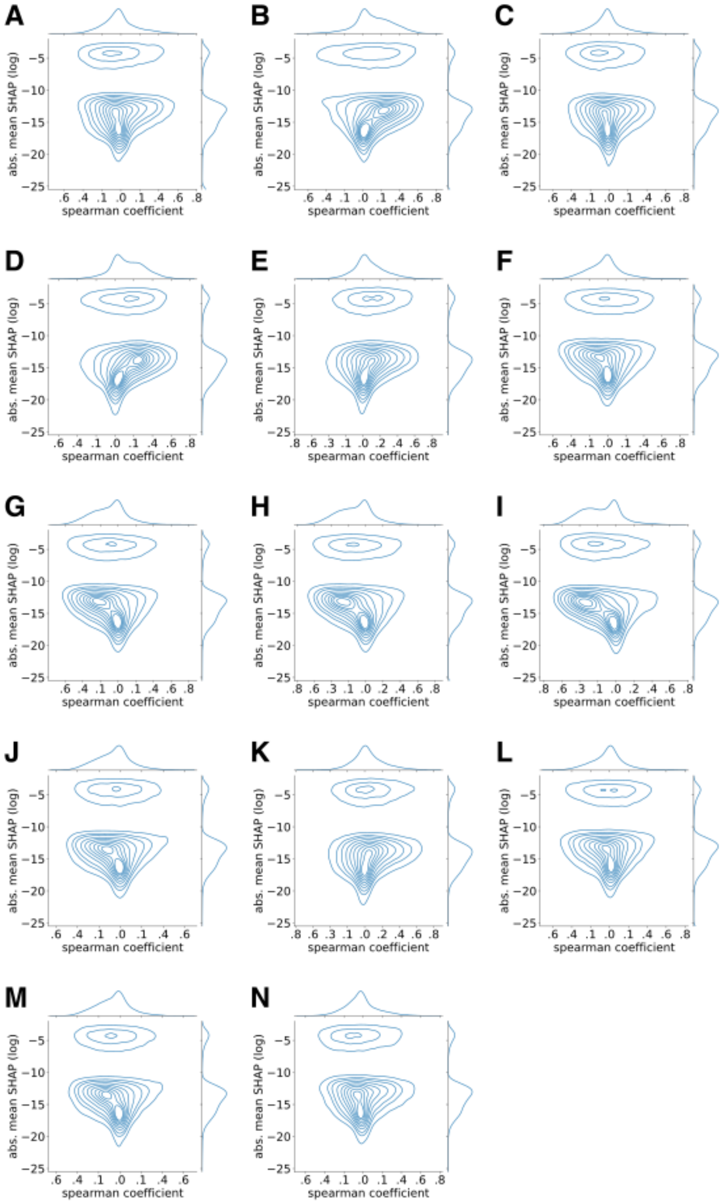


Supplemental Figure 3: **Model interpretation evaluating transcript impact on specific protein predictions.** SHAP analysis was performed for several of the most well-predicted proteins using all samples in the dataset. A kernel density graph (KDE) was plotted for mean absolute SHAP (y axis) versus spearman coefficient (x axis) for each transcript's relation to (A) CAVIN1, (B) FERMT2, (C) FLNA, (D) HCLS1, (E) HK3, (F) MCM3, (G) MCM4, (H) MCM6, (I) MMP14, (J) P4HB, (K) PTPN6, (L) SMC2, (M) STAT1, and (N) VCL proteins specifically.

| | direct | correlated | uncorrelated | noCount |
|---|---|---|---|---|
| **IG_V_gene** | 69 | 62 | 14 | 0 |
| **IG_V_pseudogene** | 0 | 26 | 149 | 8 |
| **TEC** | 0 | 739 | 271 | 6 |
| **TR_V_gene** | 0 | 83 | 23 | 0 |
| **lncRNA** | 0 | 10851 | 5539 | 112 |
| **miRNA** | 0 | 8 | 497 | 668 |
| **misc_RNA** | 0 | 386 | 641 | 11 |
| **processed_pseudogene** | 0 | 4275 | 5760 | 144 |
| **protein_coding** | 7743 | 10700 | 1438 | 18 |
| **rRNA_pseudogene** | 0 | 6 | 370 | 113 |
| **snRNA** | 0 | 68 | 1186 | 409 |
| **snoRNA** | 0 | 164 | 110 | 54 |
| **transcribed_processed_pseudogene** | 0 | 352 | 145 | 4 |
| **transcribed_unitary_pseudogene** | 0 | 114 | 41 | 0 |
| **transcribed_unprocessed_pseudogene** | 0 | 654 | 254 | 7 |
| **unprocessed_pseudogene** | 0 | 649 | 1820 | 78 |

Supplemental Table 1: **Biological function categorization of transcripts for each experimental predictive category.**
